# Genetic depletion in zebrafish uncovers requirement for septins in haematopoiesis

**DOI:** 10.64898/2026.05.05.722915

**Authors:** Kathryn Wright, Hannah Painter, Naisha Sachdev, Alexandra Budnikova, Lucy Copper, Rui Monteiro, Serge Mostowy

## Abstract

Haematopoiesis and differentiation of immune cells from haematopoietic stem and progenitor cells (HSPCs) are essential to core aspects of health and disease. A key player in haematopoiesis and HSPC differentiation is the cytoskeleton, which governs cell division and lineage bias. Despite insights using mouse models, regulation of haematopoiesis by the septin cytoskeleton is mostly unknown. Septins are unconventional filament forming proteins best known for roles in cell division and host defence. To investigate septin-mediated host defence *in vivo*, we generated septin-deficient zebrafish models for infection with *Mycobacterium marinum*. Unexpectedly, septin-deficient larvae were protected from mycobacterial infection due to significantly increased macrophage numbers, reduced cell death, and enhanced inflammatory responses. Underlying this, we found that septin-deficient larvae produce significantly more HSPCs and show myeloid lineage bias, establishing a requirement for septins in haematopoiesis. In agreement with classical HSPC hierarchy, increased myeloid production in septin-deficient larvae is at the expense of erythroid lineage production. Our findings that septins play a role in haematopoiesis is consistent with hallmarks of haematological disorders in which septin dysfunction has been implicated, including acute myeloid leukaemia, myelodysplastic syndrome, and platelet disorder Bernard-Soulier syndrome. These results highlight zebrafish as a new model to investigate septin-mediated haematopoiesis and application of septin-based medicines to treat blood disorders.

## Introduction

Haematopoiesis and haematopoietic stem and progenitor cell (HSPC) differentiation underlie fundamental aspects of immunity, from clearing invading pathogens to responding to inflammatory and neoplastic disorders^1^. HSPCs are self-renewing cells which progress to mature blood and immune cells, a process which is tightly controlled and highly conserved across species^2^. There is an emerging role for the cytoskeleton in regulation of HSPC egress and release into circulation, and in lineage bias^3–8^. Actin, microtubules, and intermediate filaments have also been shown to modulate HSPC differentiation, through interactions with RNA polymerases^9^, transcription factors involved in cell differentiation^10,11^, and contributing to nuclear morphology and deformation^12–14^.

A fourth component of the cytoskeleton are septins, discovered in yeast (*Saccharomyces cerevisiae*) as essential for cell division^15,16^. Since their discovery, septins have been associated to a wide variety of human infectious and inflammatory diseases, as well as blood disorders (including acute myeloid leukaemia [AML] and myelodysplastic syndrome [MDS]) and platelet syndromes^17–21^. Septins are GTP-binding proteins highly conserved in vertebrates (including humans and zebrafish) and classified into four groups based on sequence homology^16,22,23^ (**Supp Fig 1A**). Members from each of the four septin groups oligomerise to form complexes and higher-order structures, including filaments and ring-like structures, to carry out their physiological function (**Supp Fig 1B**). While their role in septum formation during cell division is well characterised in *S. cerevisiae* and animal tissue culture cells^16^, the breadth of their physiological functions remains poorly understood.

Despite recent insights into cytoskeletal control of haematopoiesis by actin^4,7,8^ and microtubules^3,5^, the contribution of septins is mostly unknown. Considering their interactions with actin and microtubules^24,25^, and their involvement in multiple blood disorders^18,21^, it is of great interest to explore septin roles in haematopoiesis and HSPC differentiation. Studies using mouse and pluripotent stem cell models of septin deficiency have suggested that septins are important for haematopoiesis^17,19,20,26-30^. For example, in *in vitro* cell culture and *in vivo* mouse models, loss of Septin 6 (Sept6) impacts HSPC engraftment and differentiation within lymphoid lineages, as well as cell cycle and myelopoiesis^20,27^, while loss of Septin 9 (Sept9) in T lymphocytes impacts development, overall numbers, and motility^30,31^. Considering these fundamental roles in cell production, motility, and lineage bias, there is pressing need to investigate septin functions in haematopoiesis and downstream immunity.

Zebrafish (*Danio rerio*) are a powerful vertebrate model, owing to the high level of conservation with human development and extensive genetic and transgenic tools available, providing landmark insights into haematopoiesis and blood disorders with direct impact on human health^32,33^. Moreover, previous work using zebrafish has shown that septins control inflammation and host immune responses to important human pathogens, including the World Health Organisation high priority pathogen *Shigella flexneri*^34–36^. In this study, we questioned whether septins act upstream of immune cell responses during haematopoiesis to promote inflammation and infection control. We used zebrafish infection to visualise how septin deficiency impacts haematopoiesis and immunity against *Mycobacterium marinum*, a natural fish pathogen which models *M. tuberculosis* infection of humans^37^. Using both septin mutant and knockdown zebrafish larvae, we discover that loss of septins promotes control of mycobacterial infection through increased macrophage production and enhanced inflammatory responses. Strikingly, septin-deficient larvae produce significantly more HSPCs during early haematopoiesis. We demonstrate that differentiation and lineage preferences of septin-deficient HSPCs are biased towards macrophage production at the expense of erythroid lineage production. We further show that septin-dependent haematopoietic defects can be rescued through pharmacological treatment, solidifying septins as a druggable target and highlighting the potential of septin-based medicines. Together, our work highlights unexpected roles for the cytoskeleton in haematopoiesis and identifies septins as key determinants of lineage bias.

## Materials and methods

### Ethics statement

Animal experiments were performed according to the Animals (Scientific Procedures) Act 1986 and approved by the Home Office Project license PPL PP5900632.

### Zebrafish husbandry

Adult zebrafish were housed in the Biological Services Facility at LSHTM in a 14/10-hour light/dark cycle, and larvae obtained by natural spawning were maintained at 28.5°C in E3 medium. When required (e.g. for whole mount *in situ* hybridisation and haemoglobin staining), embryos were maintained in E3 + 1-phenyl-2-thiourea (PTU) (0.036 g/L) from 1 day post fertilisation (dpf) to prevent pigmentation.

The following transgenic zebrafish lines were used: to label macrophages: *Tg(mpeg1::Gal4-FF)*^gl25^/*Tg(UAS:LIFEACT-GFP)*^mu271^,^38^ and *Tg(mpeg1::Gal4-FF)*^gl25^/*Tg(UAS-E1b::nfsB*.*mCherry)*^c264^,^39^. To label neutrophils: *Tg(lyzC:DsRed2)*^*nz50*^,^4*0*^ *Tg(lyz:nfsB-mCherry)*^*sh260*^,^41^ *and Tg(mpx:GFP)*^*ill4*^,^42^. To label HSPCs: *Tg(runx1P2:Citrine)*^*ox1Tg* 4*3*^ and *Tg(Mmu*.*Runx1:NLS-mCherry)*^*cz2010* 44^. *Tg(−6*.*0itga2b:EGFP)*^*la2Tg*^ (referred to as *cd41:GFP*) for thrombocytes (GFP^high^) and HSPCs (GFP^low^)^45^, and *TgBAC(tnfa:GFP)*^*pd1028*^ to label *tnfa*^4*6*^.

Mutant zebrafish used were *sept6* WT and *sept6*^*ls01*^ null mutants^47^, which possess a 5bp deletion in the GTP-binding domain of Sept6, and *sept15* WT and *sept15*^*sa44249*^ null N-ethyl-N-nitrosourea (ENU) mutants^47^, which harbour a single nucleotide change resulting in a premature stop codon (**Supp Fig 1C**).

### Artificial Intelligence (AI) based phenotyping

#### Dataset

A dataset consisting of approximately 500 images of embryos per line acquired at 24 hours post fertilisation (hpf) was randomly partitioned into training (90%) and test (10%) subsets. To increase the size of the training dataset, images were randomly augmented during each epoch by applying horizontal or vertical flips, or by rotating the image by 20 degrees (each with 50% probability). Images were resized to 224 × 224 pixels. Embryo detection was performed using object-detection algorithms taken from EmbryoNet^48^.

#### Classification

The classifier used was based on a convolutional neural network architecture adapted from EmbryoNet^48^, modified for 2 classes, and adapted for the Zeiss image format (czi). Models were trained using the Adam optimizer over 300 epochs with a learning rate of 10-3, a batch size of 32, using cross entropy loss. Classification accuracy was determined by testing the model on an unseen test dataset and comparing predicted and ground truth labels. Classifier and example training data are available at https://github.com/Sasha-b-code/Sept6_Classifier

#### Zebrafish injections

For CRISPR-Cas9 mediated knockdown, embryos were injected at the one- to two-cell stage with 1 nL of CRISPR mixture containing 1 μg/μL pooled guide RNA (gRNA) (**Table 1**) and 500 μg/mL Cas9. gRNA was synthesised as previously described^49^. Primers containing gene-specific guide sites and a T7 promotor were annealed to a standard scaffold primer. gRNA was *in vitro* transcribed from purified DNA templates (HiScribe T7 High Yield RNA Synthesis Kit, NEB), and purified products used for injection. Embryos were injected with either a gene-specific gRNA or a scramble control gRNA.

**Table 1.**
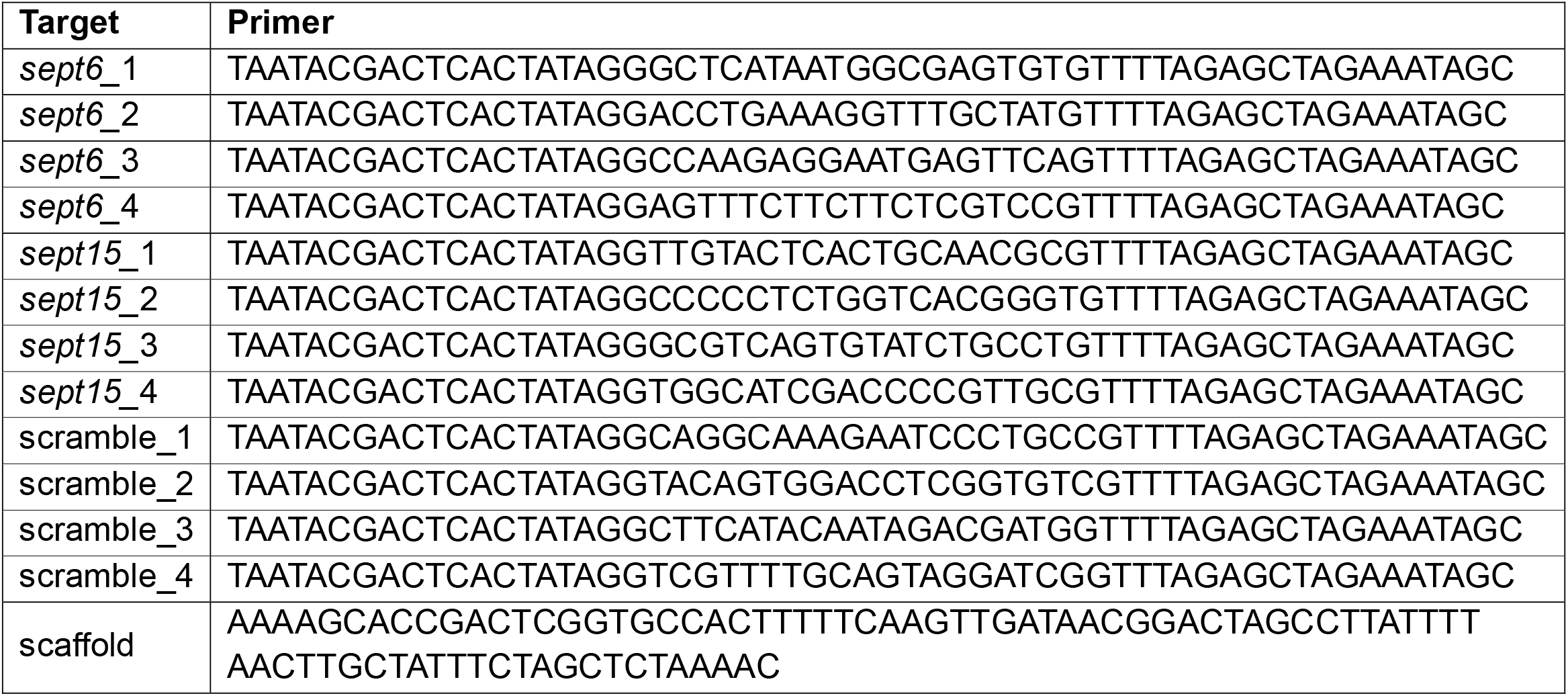
Guide RNA sequences used for CRISPR-Cas9 knockdowns.

For bacterial infections, larvae were anaesthetised with 160 μg/mL of Tricaine (MS-222) and infected via the caudal vein with ~200 CFU of *M. marinum* M strain, either unlabelled WT or expressing Wasabi, tdTomato, or Katushka fluorescent proteins. Bacterial culture was performed as previously described^50,51^. Briefly, *M. marinum* was grown at 33°C in 7H9 supplemented with OADC and 50 μg/mL Hygromycin (for fluorescent bacteria) to an OD600 of 0.6-0.8. Single cell suspensions were prepared by repeated washing, shearing, and aspiration through a 28 G needle, prior to filtration through a 5 μm filter. Aliquots of enumerated bacteria were frozen and stored at −80°C and diluted with 0.5% phenol red immediately prior to injection. Following infection, larvae were recovered in fresh E3 media with or without PTU and maintained at 28.5°C.

### Macrophage depletion

Transgenic macrophage zebrafish were treated with 10 mM metronidazole (MTZ) or DMSO vehicle control in E3 media for 18 hrs at 28.5°C protected from light.

### Forchlorfenuron treatment

Larvae were randomly assigned to treatment groups and treated with either 25 μM of the small molecule forchlorfenuron (FCF) or DMSO vehicle control from 4 hpf and refreshed daily.

### Whole mount *in situ* hybridisation

Whole mount *in situ* hybridisation (WISH) was carried out as previously described^52^ using probes for *runx1, cmyb, rag1*, and *hbbe1*. 36 hpf or 4 dpf embryos were subjected to WISH and imaged using a Nikon SMN800N stereomicroscope and a MOTIC S6 camera. Quantification of *runx1, cmyb*, and *rag1* staining was performed by converting images to grayscale and inverted using Fiji. Pixel intensity was measured for the appropriate staining area relative to an unstained background to obtain pixel intensity values for each embryo and probe^53^. For analysis of *hbbe1*, staining was categorised by localisation: no peripheral staining (i.e. heart only), caudal haematopoietic tissue only (CHT only), or CHT and circulatory loop (CHT and posterior cardinal vein/dorsal aorta [PCV/DA]).

### Whole mount staining

Larvae were infected with *M. marinum* via caudal vein injection and analysed for and apoptotic cells at 3 dpi using the Click-iT Plus TUNEL Assay (Thermo Fisher Scientific) as previously described^51^. Larvae were fixed in 4% PFA overnight prior to permeabilisation. edUTP and Alexa Fluor 647 incorporation reactions were performed at 37°C protected from light, and stained larvae stored in 50% glycerol until imaging.

For detection of haemoglobin in erythrocytes, uninfected larvae were collected at required timepoints and subjected to *o-*dianisidine staining^54^. Larvae were fixed in 4% PFA overnight and washed in PBST (PBS + 0.1% Tween-20). Embryos were stained with *o-*dianisidine working solution (0.6 mg/mL *o*-dianisidine, 0.01 M sodium acetate, 0.65% H_2_O_2_, 40% vol/vol ethanol) for 30 minutes in the dark, followed by PBST washes and postfixing in 4% PFA overnight. Embryos were transferred to 50% glycerol prior to imaging.

### Cellular reactive oxygen species staining

For reactive oxygen species (ROS) detection, 3 dpi *M. marinum* infected larvae were incubated in 5 μM CellROX Green (Thermo Fisher Scientific) for 30 minutes at 28.5°C and imaged immediately.

### Microscopy and image analysis

Live imaging of larvae was performed on anaesthetised embryos on a depression microscope slide, or within a 96-well black-walled microplate (PerkinElmer), using either a Leica M205FA Fluorescent Stereo Microscope equipped with a Leica DFC365FX monochrome digital camera, or a Zeiss Cell Discoverer 7.

Bacterial burden and fluorescent cell area were analysed using Fiji to quantify the fluorescent pixel count, defined as fluorescent signal above a consistent set background determined empirically for each experimental dataset^37,50^. Data are presented as total fluorescent area (pixels) above background level. To analyse infection-associated *tnfa*, expression of GFP in the *TgBAC(tnfa:GFP*) line was measured within a 500 μm box around infection foci, and for sterile inflammation tail wound assays, cells and *tnfa* expression within 100 μm of the tail resection site were analysed, as previously described^51,55^.

## Statistical analysis

Statistical analysis was performed in GraphPad Prism (v.11.0.0) using default parameters. Non-parametric data was analysed by a Mann–Whitney test or Kruskal– Wallis test depending on experimental design, and comparisons between groups performed using corrected Dunn’s test. Normally distributed data were analysed with two-way ANOVA and Sidak multiple comparisons test as indicated in figure legends. Outliers were removed before statistical analysis using ROUT, with Q = 1%.

## Results

### Septin-deficient larvae control mycobacterial infection

Our previous work combining septin morphants and mutants established a role for Septin 2 (*sept2*) and Septin 15 (*sept15*) in host response to infection^34–36,47^, with *sept2*- and *sept15*-depleted larvae displaying increased susceptibility to *S. flexneri*. To further explore the role of septins in host defence, we infected Septin 6 (*sept6*)-deficient larvae with *M. marinum*. Septin 6 is a core member of septin complexes, essential for their cooperative function^56^. We recently generated a *sept6* mutant zebrafish line, which does not exhibit any developmental defects or transcriptional compensation from other septin genes^47^. In agreement, an AI classifier trained on both WT and *sept6*-deficient larvae to identify phenotypic and developmental defects failed to reliably separate the two groups (**Supp Fig 1D**). Strikingly, we observed that *sept6*-deficient larvae better control mycobacterial infection as compared to controls, significantly reducing bacterial load (1.8-fold) by 3 days post infection (dpi) (**Fig 1A**). These results are in contrast to what has been observed with the inflammatory pathogen *S. flexneri*^47^. A hallmark of septin biology is oligomerisation of subunits from each of the four groups into complexes^16^; we therefore queried whether control of *M. marinum* burden was conserved across septin groups. To test this, we infected *sept15* (zebrafish homolog of human SEPT7 and central subunit in complexes^56^) - deficient zebrafish with *M. marinum* and observed the same protective effect as for *sept6*-deficient larvae (**Supp Fig 2A**), suggesting that septin roles in host defence are as an oligomeric complex.

**Figure 1.**
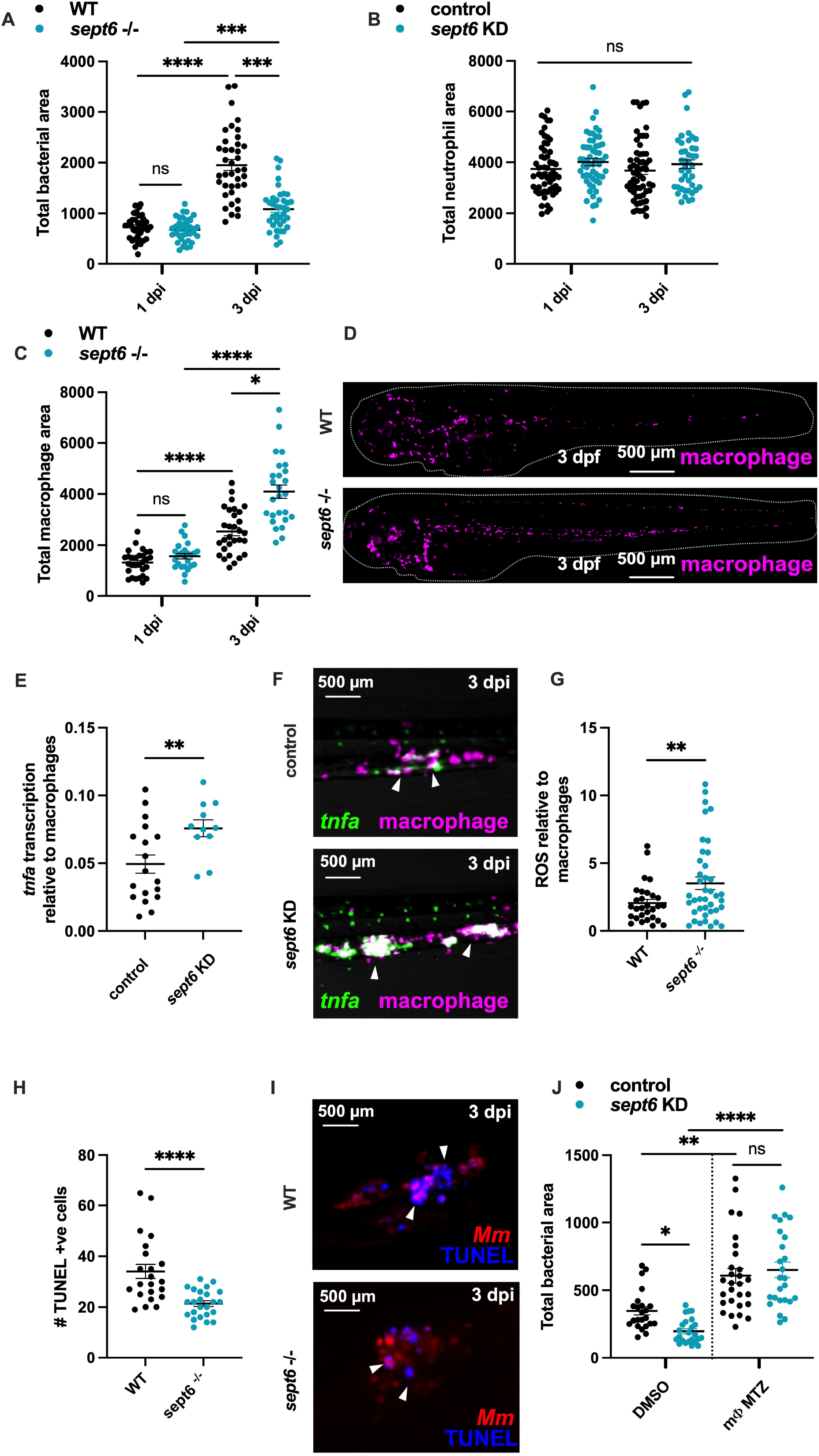
Septin 6 directs macrophage inflammatory responses during mycobacterial infection. **(A)** *M. marinum* bacterial burden in WT and *sept6*^*−/−*^ larvae at 1 and 3 dpi. **(B)** Quantification of whole-body neutrophil levels in *M*. marinum-infected scramble control and *sept6* knockdown [KD] larvae. **(C)** Quantification of whole-body macrophage levels in *M. marinum*-infected WT and *sept6*^*−/−*^ larvae. **(D)** Representative images of macrophages in *Tg(mpeg1::Gal4-FF)*/*Tg(UAS:LIFEACT-GFP)* WT and *sept6*^*−/−*^ larvae at 3 dpf **(E)** Measurement of *tnfa* promotor activation relative to total macrophage levels at 3 dpi following infection with *M. marinum* in scramble control and *sept6* knockdown [KD] larvae. **(F)** Representative images of *tnfa* promoter-driven GFP expression in macrophages (magenta) using the double transgenic *TgBAC(tnfa:GFP); Tg(m peg1::Gal4-FF)*/*Tg(UAS:LIFEACT-GFP)* line at 3 dpi in *M. marinum* infected scramble control and *sept6* knockdown [KD] larvae. **(G)** Quantification of cellular ROS relative to total macrophage levels at 3 dpi in *M. marinum-*infected WT and *sept6*^*−/−*^ larvae stained with CellROX. **(H)** TUNEL +ve cells in *M. marinum*-infected WT and *sept6*^*−/−*^ larvae were counted at 3 dpi. **(I)** Representative images of TUNEL staining at 3 dpi in WT and *sept6*^*−/−*^ larvae. TUNEL +ve cells are blue and *M. marinum* is red. **(J)** *M. marinum* burden in control and macrophage-depleted (mΦ MTZ) scramble control and *sept6* knockdown [KD] larvae at 3 dpi. Each data point represents a single measurement (larvae) with the mean and SEM shown. Graphs are representative of three experimental replicates with the exception of ROS measurements, which was performed in duplicate. * P ≤ 0.05, ** P ≤ 0.01, *** P ≤ 0.001, **** P ≤ 0.0001.

Considering that septins control inflammation^34,57^, we hypothesised that outcome of infection in septin-deficient larvae is dependent on the pathogen. For example, restriction of inflammation may enhance control of inflammatory pathogens such as *S. flexneri*, while chronic infections caused by *M. marinum* may benefit from restriction of inflammation. We first looked at neutrophils, one of the earliest cells to respond to infection and inflammation but did not observe any differences in neutrophil numbers between WT and *sept6*-deficient larvae at either 1 or 3 dpi (**Fig 1B**). We next assessed macrophages, as they are widely recognised in zebrafish and humans as crucial for mycobacterial control^58,59^. Surprisingly, we noted that *sept6*-deficient larvae had a greater number of macrophages at 3 dpi (1.6-fold; **Fig 1C-D**); the same was observed for *sept15-*deficient larvae (**Supp Fig 2B**). To determine whether these macrophages were more responsive to *M. marinum*, and thus contributing to reduced bacterial burden, we assessed infection-induced inflammation. We performed a CRISPR-Cas9-mediated knockdown of *sept6* in the transgenic *TgBAC(tnfa:GFP)*^*pd1028*^ line, to measure amount of *tnfa* promotor activation at infection foci. Consistent with previous work showing that septins restrict inflammation during *S. flexneri* infection^34^, there was enhanced *tnfa* production in *sept6*-deficient larvae at 3 dpi (**Supp Fig 2C-D**). The increase in inflammatory *tnfa* was still observed when normalised to macrophage levels to account for the septin-dependant increase in macrophage production (**Fig 1E-F**). Further, at 3 dpi an increase in cellular ROS production (important inflammatory and antibacterial product involved in host response to mycobacterial infection) was increased in *sept6-* deficient larvae when normalised to macrophage numbers, suggesting that *sept6*-deficient macrophages produce more ROS than WT cells (**Fig 1G**). Despite an increase in infection-associated *tnfa* promotor activation, there was no difference in number of *tnfa*-expressing neutrophils at infection foci (**Supp Fig 3A-D**), suggesting that increased macrophages are strictly responsible for *tnfa* production. In agreement with reduced cell death being important for bacterial control, as well as macrophage aggregation and granuloma formation during early infection^60,61^, we assessed apoptotic cell death and detected a significant decrease in TUNEL-stained cells in 3 dpi *M. marinum* infected larvae (**Fig 1H-I**).

To confirm that inflammatory macrophages in *sept6-*deficient larvae were controlling infection, we depleted macrophages prior to infection and nullified the protective effect of *sept6-*deficiency (**Fig 1J**). Together, we conclude that the greater number of macrophages and their enhanced inflammatory response are cooperatively restricting infection and mycobacterial burden in septin-deficient larvae.

### Septin deficiency enhances macrophage inflammatory potential

Our results showed that *sept6*-deficient macrophages were more inflammatory following infection with *M. marinum*; we next queried whether this was specific to infection or a broad response to stimulation. To test this, we induced sterile inflammation using a zebrafish tail resection model^62^. Resection of the zebrafish tail at the pigment gap is distal to the circulatory loop and sufficient to induce chemotaxis and immune cell migration^63^. In this case, *sept6*-deficient larvae displayed a significant decrease in macrophage recruitment to the wound site at both 3- and 6-hours post wounding (hpw) (2.1-fold and 2.5-fold respectively) (**Fig 2A-B**), however recruited macrophages were more inflammatory with a greater number expressing *tnfa* (**Fig 2C-D**). Consistently, *sept15*-deficient macrophages showed a significant decrease (1.4-fold) in recruitment to the wound site with significantly increased expression of *tnfa* (**Supp Fig 4A-B**).

**Figure 2.**
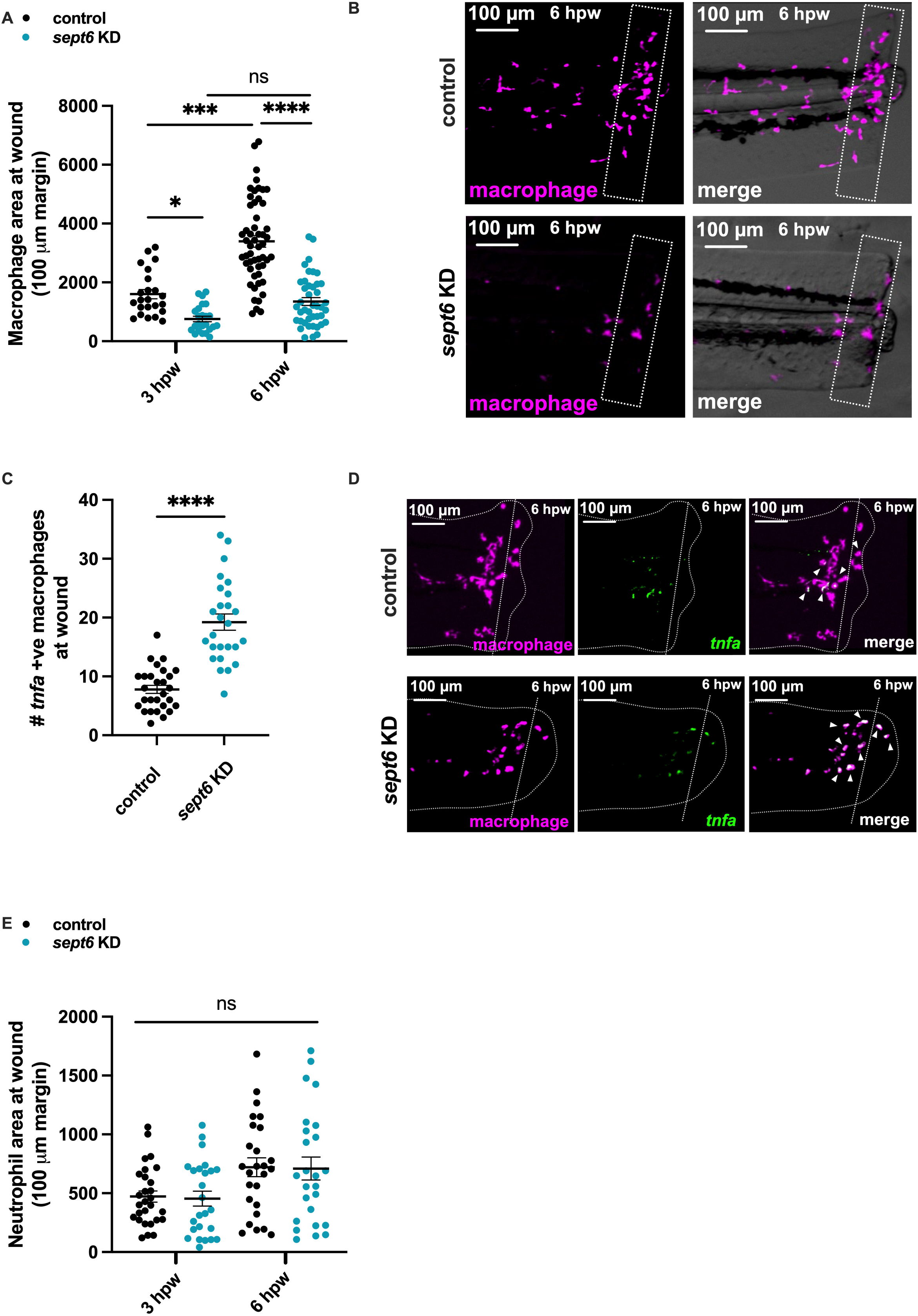
*sept6*-deficient macrophages exhibit enhanced inflammatory potential. **(A)** Measurement of macrophage recruitment to a tail wound in WT and *sept6* knockdown [KD] larvae at 3 and 6 hpw. **(B)** Representative images of macrophage (magenta) recruitment to tail wound sites in *Tg(mpeg1::Gal4-FF)*/*Tg(UAS:LIFEACT-GFP)* scramble control and *sept6* knockdown [KD] larvae at 6 hpw. Dotted rectangles indicate the region of measurement (100 μm from wound site). **(C)** Quantification of macrophage-specific *tnfa* promotor activation at the wound site in scramble control and *sept6* knockdown [KD] larvae at 6 hpw. **(D)** Representative images of *tnfa* promoter-driven GFP expression in macrophages (magenta) at the wound site in double transgenic *TgBAC(tnfa:GFP); Tg(mpeg1::Gal4-FF)*/*Tg(UAS:LIFEACT-GFP)* scramble control and *sept6* knockdown [KD] larvae at 6 hpw. Dotted line indicates tail wound cut site. **(E)** Quantification of neutrophil recruitment to a tail wound in scramble control and *sept6* knockdown [KD] larvae at 3 and 6 hpw. Each data point represents a single measurement (larvae) with the mean and SEM shown. Graphs are representative of three experimental replicates. * P ≤ 0.05, ** P ≤ 0.01, *** P ≤ 0.001, **** P ≤ 0.0001.

Similar to results using *M. marinum* infection, we observed that *sept6*-dependent inflammatory defects do not impact neutrophils, with no difference in neutrophil number or inflammatory state observed between WT and *sept6*-deficient larvae (**Fig 2E**). Together, we conclude that enhanced inflammatory potential of *sept6*-deficiency is specific to macrophages, and not a universal feature of myeloid populations. Further, the septin-dependent increase in macrophage numbers and inflammatory response is conserved across different septin groups, indicating that these effects are due to disruption of septin hetero-oligomeric complexes.

### Septin deficiency promotes haematopoiesis and macrophage production

We considered that the greater number of macrophages in *M. marinum-*infected *sept6*-deficient larvae was because of increased inflammation and reduced cell death. To test this, we analysed macrophages in uninfected larvae from 28 hours post fertilisation (hpf). Surprisingly, we discovered that increased macrophage production in *sept6-*deficient larvae begins between 28-48 hpf (**Fig 3A**), prior to time of infection (48-54 hpf). An increase in macrophages was also observed in uninfected *sept15*-deficient larvae at 3 dpf (**Supp Fig 4C**). This shows that macrophage increase in *sept6-*deficient larvae is independent of infection and likely due to myelopoietic bias (rather than reduced cell death).

**Figure 3.**
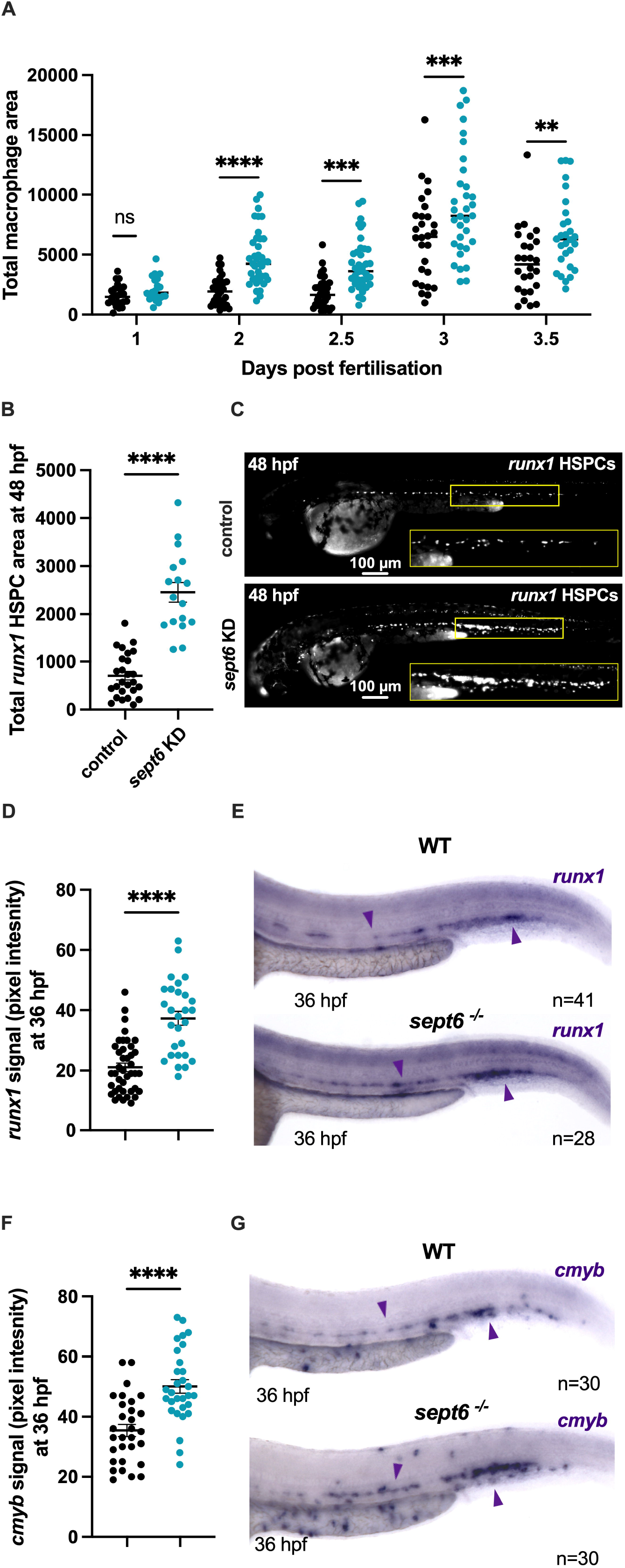
Loss of *sept6* increases haematopoiesis and drives macrophage production. **(A)** Quantification of whole-body macrophage levels in uninfected WT and *sept6*^*−/−*^ larvae across development from 1-3.5 dpf. **(B)** Quantification of *runx1* HSPC fluorescent area at 48 hpf in WT and *sept6* knockdown [KD] larvae. **(C)** Representative images of *runx1* HSPCs at 48 hpf in *Tg(runx1P2:Citrine)* scramble control and *sept6* knockdown [KD] larvae. Insets show the caudal haematopoietic tissue region and residing HSPCs. **(D)** Quantification of WISH staining for *runx1* in 36 hpf WT and *sept6*^*−/−*^ larvae **(E)** Representative images of WISH staining for *runx1* in WT and *sept6*^*−/−*^ larvae at 36 hpf. **(F)** Quantification of WISH staining for *cmyb* in 36 hpf WT and *sept6*^*−/−*^ larvae **(G)** Representative images of WISH staining for *cmyb* in WT and *sept6*^*−/−*^ larvae at 36 hpf Each data point represents a single measurement (larvae) with the mean and SEM shown. Graphs are representative of three experimental replicates with the exception of WISH measurements, which was performed in a single experiment with a minimum of 20 biological replicates. * P ≤ 0.05, ** P ≤ 0.01, *** P ≤ 0.001, **** P ≤ 0.0001.

To understand at what stage septins regulate immune cell production, we assessed HSPCs in septin-deficient larvae using transgenic *runx1* zebrafish lines, where the essential haematopoietic transcription factor Runx1 labels HSPCs and hemogenic endothelium^43^. We discovered that *sept6* and *sept15* deficiency significantly increased HSPC production at 48 hpf (**Fig 3B-C, Supp Fig 4D**), suggesting that septins impact macrophage production through upstream haematopoiesis. To confirm septin-dependant haematopoietic defects, we performed whole-mount *in situ* hybridisation (WISH) of HSPC markers *runx1* and *cmyb* in *sept6*-deficient larvae at 36 hpf. Consistent with knockdown data, expression of *runx1* and *cmyb* was significantly increased (1.8- and 1.4-fold respectively) in *sept6* mutants compared to WT (**Fig 3D-G**).

To probe the relationship between HSPC and macrophage numbers in *sept6* mutants, we tested if there was a feedback loop, with macrophages signalling to HSPCs to enhance production. However, depletion of macrophages from 24 hpf did not impact HSPC numbers at 48hpf (**Supp Fig 5A-B**), highlighting that septins are important for both haematopoiesis and macrophage bias.

### Septins balance myeloid and erythroid lineage production

To test if increased production of immune cells is specific to macrophages, we depleted macrophages in *Tg(mpeg1::Gal4-FF)*/*Tg(UAS-E1b::nfsB*.*mCherry)* transgenic line at 48 hpf using metronidazole (MTZ) but observed no change in neutrophil production (**Fig 4A**). These experiments reveal that shifts in lineage bias due to septin deficiency is specific to macrophages, rather than the broader myeloid population.

**Figure 4.**
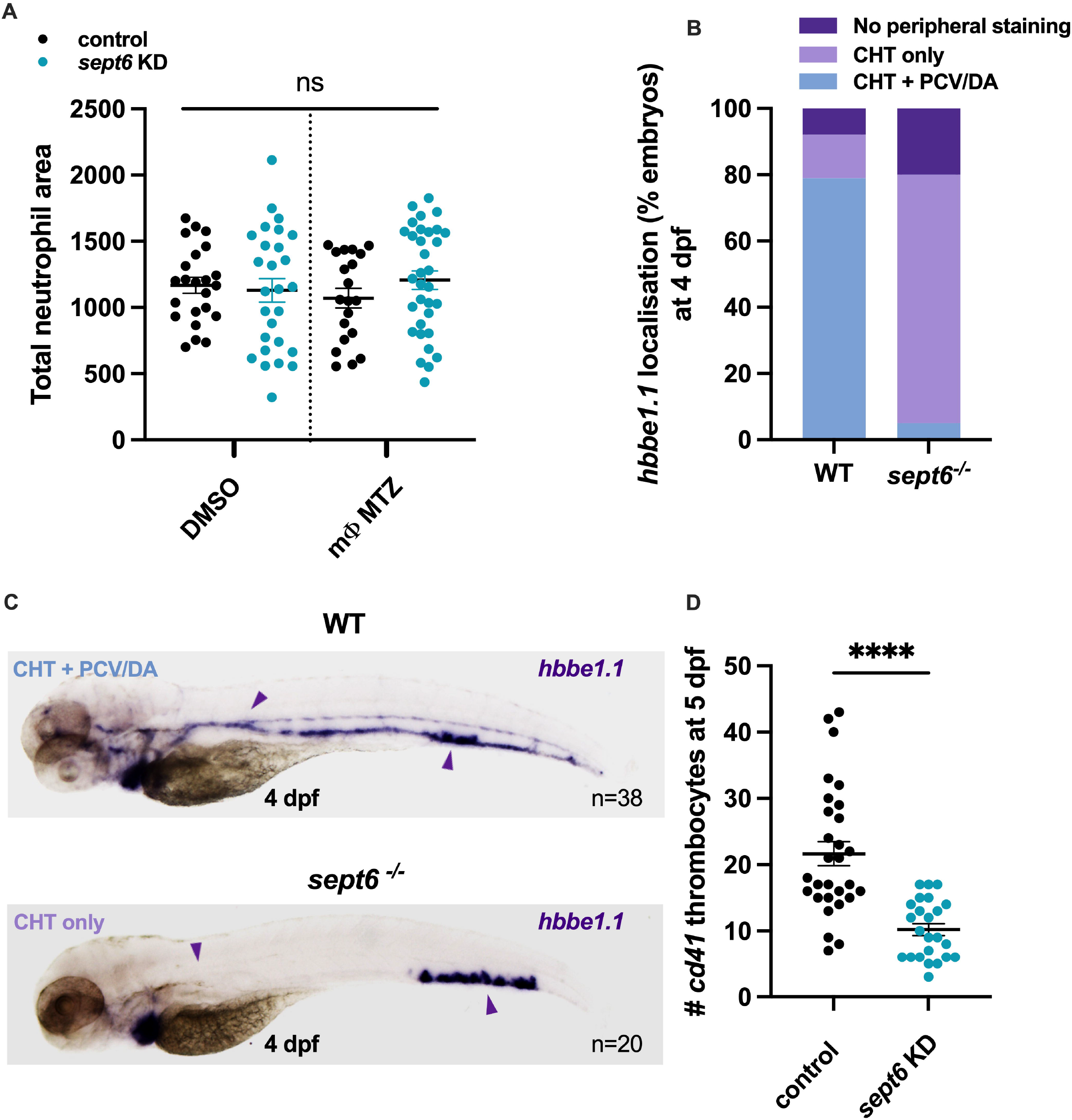
*sept6*-deficiency enhances macrophage production at the expense of erythroid lineages. **(A)** Quantification of whole-body neutrophil levels in control and macrophage-depleted (mΦ MTZ) scramble control and *sept6* knockdown [KD] larvae at 48 hpf. **(B)** Localisation of *hbbe1*.*1* WISH staining in WT and *sept6*^*−/−*^ larvae at 4 dpf. **(C)** Representative images of *hbbe1*.*1* WISH staining in 4 dpf WT and *sept6*^*−/−*^ larvae. WT larvae show staining throughout the circulation and *sept6*^*−/−*^ larvae display staining localised to the CHT region only. **(D)** Quantification of thrombocytes at 5 dpf in scramble control and *sept6* knockdown [KD] larvae. Each data point represents a single measurement (larvae) with the mean and SEM shown. Graphs are representative of three experimental replicates with the exception of WISH measurements, which was performed in a single experiment with a minimum of 20 biological replicates. * P ≤ 0.05, ** P ≤ 0.01, *** P ≤ 0.001, **** P ≤ 0.0001.

To resolve the increased macrophage production, we next evaluated multiple blood cell lineages in *sept6*-deficient larvae. Considering that HSPC-derived myeloid progenitors will differentiate to either myeloid or erythroid lineages during early haematopoiesis^64^, we assessed the erythroid lineage as well and performed WISH of the globin gene *hbbe1*.*1* in 4 dpf larvae in *sept6*-deficient larvae. Consistent with the classic myeloid versus erythroid paradigm, in which myeloid progenitors differentiate into megakaryocyte or granulocyte-monocyte lineages^65^, we observed significant differences in staining of mutant larvae (**Fig 4B-C**). In WT larvae, the majority (79%) exhibited staining throughout the circulatory loop; this included the caudal haematopoietic tissue (CHT) where progenitors and mature cells reside, as well as the posterior cardinal vein (PVC) and dorsal aorta (DA). Strikingly, 75% of *sept6*-deficient larvae exhibited staining localised only in the CHT region, strongly suggesting defects in mature circulating erythrocytes. This was also observed using *o-*dianisidine staining for erythrocytes (**Supp Fig 5C**). We next exploited the transgenic *cd41:GFP* line to quantify thrombocytes (GFP^high^ subset)^45^ and observed a significant decrease (2-fold) in cell numbers (**Fig 4D**), confirming that increased macrophage production in *sept6*-deficient larvae is at the expense of megakaryocyte and erythroid lineage production.

In the case of larval zebrafish, lymphocyte progenitors reside in the thymus from 3 dpf until adaptive immunity becomes functional at 3 weeks post fertilisation^66^. WISH of the T cell marker *rag1* to probe lymphocyte progenitors showed no difference between WT and Sept6 mutants at this developmental stage (**Supp Fig 5D-E**). These data suggest that septin-dependent haematopoietic defects are restricted to myeloid lineages.

Considering the emergence of septin-based medicines for human health impact^36,67,68^, we investigated their ability to correct the lineage bias in *sept6-* deficient larvae. We employed forchlorfenuron (FCF), a small molecule which alters septin assembly dynamics and stabilises complexes without impacting actin and microtubule networks^69^. Treatment with FCF reduced both *runx1*-lablled haemogenic endothelium (HE; the precursor from which HSPCs arise during epithelial to haematopoietic transition) and macrophage numbers in *sept6*-deficient larvae, highlighting the potential to correct for septin-dependant lineage bias through pharmacological treatment (**Fig 5A-D**).

**Figure 5.**
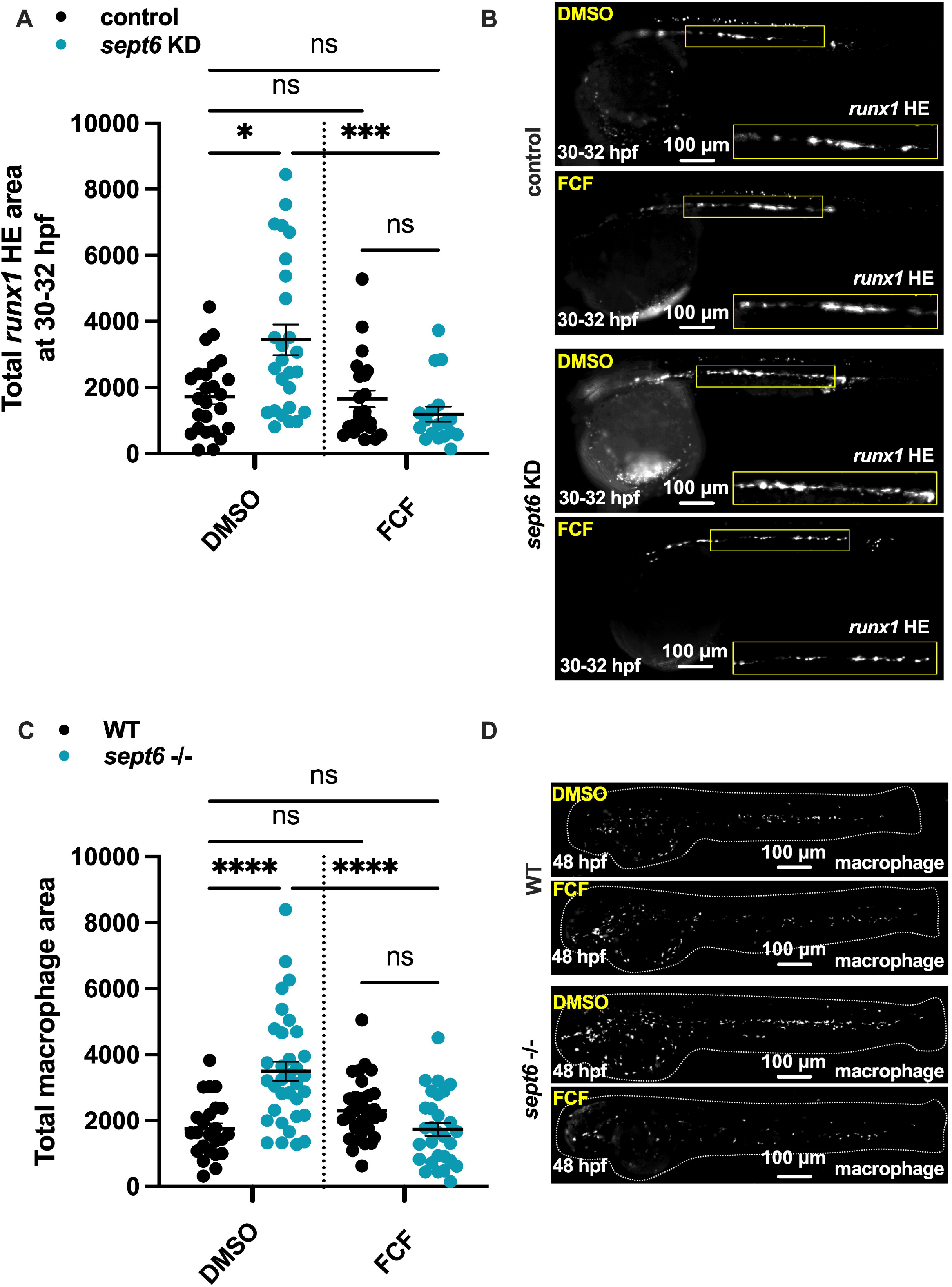
*sept6*-dependent haematopoietic defects can be rescued by forchlorfenuron. **(A)** Quantification of *runx1-*labelled HE levels in DMSO or FCF treated scramble control and *sept6* knockdown [KD] larvae at 30-32 hpf. **(B)** Representative images of *runx1*-labelled HE at 30-32 hpf in *Tg(runx1P2:Citrine)* scramble control and *sept6* knockdown [KD] larvae following treatment with DMSO or FCF. Insets enlarge the HE region. **(C)** Whole-body macrophage levels in DMSO or FCF treated WT and *sept6*^*−/−*^ larvae at 48 hpf. **(D)** Representative images of macrophages in *Tg(mpeg1::Gal4-FF)*/*Tg(UAS:LIFEACT-GFP)* WT and *sept6*^*−/−*^ larvae at 48 hpf following treatment with DMSO or FCF. Each data point represents a single measurement (larvae) with the mean and SEM shown. Graphs are representative of three experimental replicates. * P ≤ 0.05, ** P ≤ 0.01, *** P ≤ 0.001, **** P ≤ 0.0001.

## Discussion

Here we discover a fundamental role for septins in regulating haematopoiesis to drive macrophage production and host immunity. In searching to understand how septins contribute to macrophage production, we discovered that septins govern haematopoiesis, with septin-deficiency resulting in increased HSPC numbers and lineage bias and show that septin-targeting small molecules can correct dysregulated haematopoiesis. These results highlight zebrafish as a new model to investigate septin-mediated haematopoiesis and application of septin-based medicines to treat blood disorders.

By testing different septin groups, we show that septin depletion leads to an increase in HSPC numbers and lineage bias. While orphan roles for individual septin subunits have been proposed^70,71^, we conclude that septins are functioning as hetero-oligomeric complexes (rather than individually) to regulate haematopoiesis. It is next of great interest to explore how different septin complexes (e.g. hexamer vs octamer) may contribute to HSPC generation, differentiation, and cell fate.

We reveal how septin-mediated haematopoiesis and enhanced inflammatory responses impact immunity to mycobacterial infection. Previous work has discovered that septins control *Shigella* infection through regulating host inflammatory pathways and macrophage responses^34–36^, with septin-deficient larvae displaying reduced infection control. This is in contrast to our observations here with *M. marinum*, which we attribute to the distinct intracellular life cycles of the different pathogens. Future investigation of adult zebrafish, where adaptive immunity is fully functional, will be required to understand how lymphoid lineages are impacted by septin-mediated haematopoiesis, and investigate how septin mutants may respond to chronic infections that rely on lymphocyte control.

Septins are notoriously associated with blood disorders such as AML and MDS, as well as platelet and bleeding disorders^18–21,72,73^. Our zebrafish models present a powerful system to investigate mechanistic cell biology *in vivo* and dissect septin involvement in pathways responsible for haematological disorders. As we were able to correct for septin-dependant lineage bias through pharmacological treatment (i.e. septin stabilisation), further screening of septin-targeting compounds may yield new therapeutic avenues for human health. Using zebrafish as a model for *in vivo* study of septin roles in haematopoiesis, future work may illuminate how septin defects in humans contribute to haematological disorders and host response to infection.

## Acknowledgements

We thank Mostowy lab members, in particular M. Castro-Gomes, D. Broktazky, and A.T. López-Jiménez for helpful discussions. We thank the LSHTM Biological Services Facility for the work and care of our fish stocks.

## Author Contributions

Conceptualisation: K.W. and S.M. Methodology: K.W., R.M., and S.M. Investigation and formal analysis: K.W., H.P., N.S., A.B., L.C. Editing: K.W., H.P., L.C., R.M., S.M. Writing: KW and SM.

## Figure legends

**Supplementary Figure 1. Septin conservation and zebrafish mutants. (A)** Conservation of septins in vertebrates. **(B)** Septin complex structure and higher order structures. **(C)** Generation of septin mutant zebrafish and schematic of the impact of mutations on protein structure. Septin 6 mutants lack the septin unique element [SUE] and C-terminus, while Septin 15 mutants lack the C-terminus only. Schematic adapted from Torraca et.al, 2023 [47]. **(D) (I)** AI identification of *sept6*^*−/−*^ embryos. Example images of AI classifier training data **(ii)** Percentage of correctly and incorrectly assigned larvae from entire training dataset (containing both WT and mutant images) as “*sept6*^*−/−*^ larvae”.

**Supplementary Figure 2. Septin-directed macrophage responses during infection are conserved. (A)** *M. marinum* bacterial burden at 1 and 3 dpi in WT and *sept15*^*−/−*^ larvae. **(B)** Whole-body macrophage levels in *M. marinum*-infected WT and *sept15*^*−/−*^ larvae. **(C)**Total whole-body *tnfa* promotor activity in scramble control and *sept6* knockdown [KD] larvae at 3 dpi. **(D)** Representative images of *tnfa* promotor activity (green) in *M. marinum* (unlabelled) infected *TgBAC(tnfa:GFP)* scramble control and *sept6* knockdown [KD] larvae at 3 dpi. Insets show infected areas with early nascent granulomas. Each data point represents a single measurement (larvae) with the mean and SEM shown. Graphs are representative of three experimental replicates. * P ≤ 0.05, ** P ≤ 0.01, *** P ≤ 0.001, **** P ≤ 0.0001.

**Supplementary Figure 3. Loss of septins does not impact neutrophil inflammatory potential. (A)** Infection-associated *tnfa* promotor activation in *M. marinum-*infected scramble control and *sept6* knockdown [KD] larvae at 1 dpi. **(B)** Quantification of neutrophil-specific *tnfa* promotor activation in *M. marinum*-infected scramble control and *sept6* knockdown [KD] larvae at 1 dpi. **(C)** Infection-associated *tnfa* promotor activation in *M. marinum-*infected scramble control and *sept6* knockdown [KD] larvae at 3 dpi. **(D)** Quantification of neutrophil-specific *tnfa* promotor activation in *M. marinum*-infected scramble control and *sept6* knockdown [KD] larvae at 3 dpi. Each data point represents a single measurement (larvae) with the mean and SEM shown. * P ≤ 0.05, ** P ≤ 0.01, *** P ≤ 0.001, **** P ≤ 0.0001.

**Supplementary Figure 4. Septin-dependant macrophage inflammatory and haematopoietic defects are conserved across septin groups. (A)** Macrophage recruitment to a tail wound in WT and *sept15*^*−/−*^ larvae at 3 and 6 hpw. **(B)** Quantification of macrophage-specific *tnfa* promotor activation at the wound site in scramble control and *sept15* knockdown [KD] larvae at 6 hpw. **(C)** Quantification of whole-body macrophage levels in uninfected WT and *sept15*^*−/−*^ larvae at 3 dpf. **(D)** Quantification of *runx1* HSPC fluorescent area at 48 hpf in scramble control and *sept15* knockdown [KD] larvae. Each data point represents a single measurement (larvae) with the mean and SEM shown. Graphs are representative of three experimental replicates. * P ≤ 0.05, ** P ≤ 0.01, *** P ≤ 0.001, **** P ≤ 0.0001.

**Supplementary Figure 5. Septin deficiency impacts erythroid-myeloid balance but not lymphoid progenitors in zebrafish larvae. (A)** Quantification of *runx1* +ve HSPCs in control and macrophage-depleted (mΦ MTZ) *Tg(runx1P2:Citrine)* scramble control and *sept6* knockdown [KD] larvae at 48 hpf. **(B)** Representative images of *runx1* +ve HSPCs in control and macrophage-depleted (mΦ MTZ) *Tg(runx1P2:Citrine)* scramble control and *sept6* knockdown [KD] larvae at 48 hpf. Insets show the caudal haematopoietic tissue region and residing HSPCs. **(C)** Representative images of *o-*dianisidine staining 4 dpf WT and *sept6*^*−/−*^ larvae. **(D)** Quantification of WISH staining for *rag1* in 4 dpf WT and *sept6*^*−/−*^ larvae. **(E)** Representative images of *rag1* WISH staining of 4 dpf WT and *sept6*^*−/−*^ larvae. Arrows indicate the staining region in the thymi. Each data point represents a single measurement (larvae) with the mean and SEM shown.

