## Supplementary figures and images for "Genetic depletion in zebrafish uncovers requirement for septins in haematopoiesis"

### Supplementary material

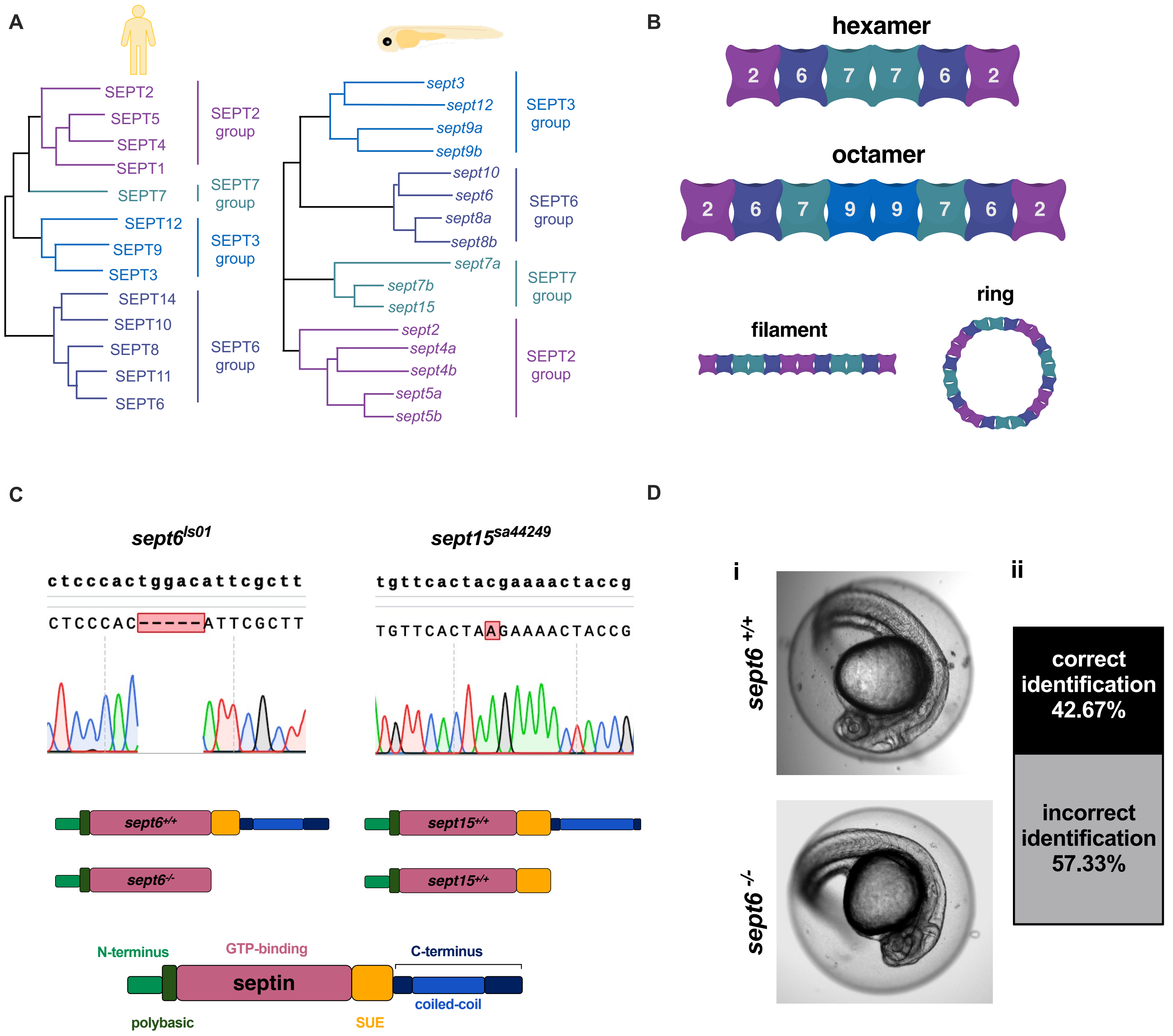

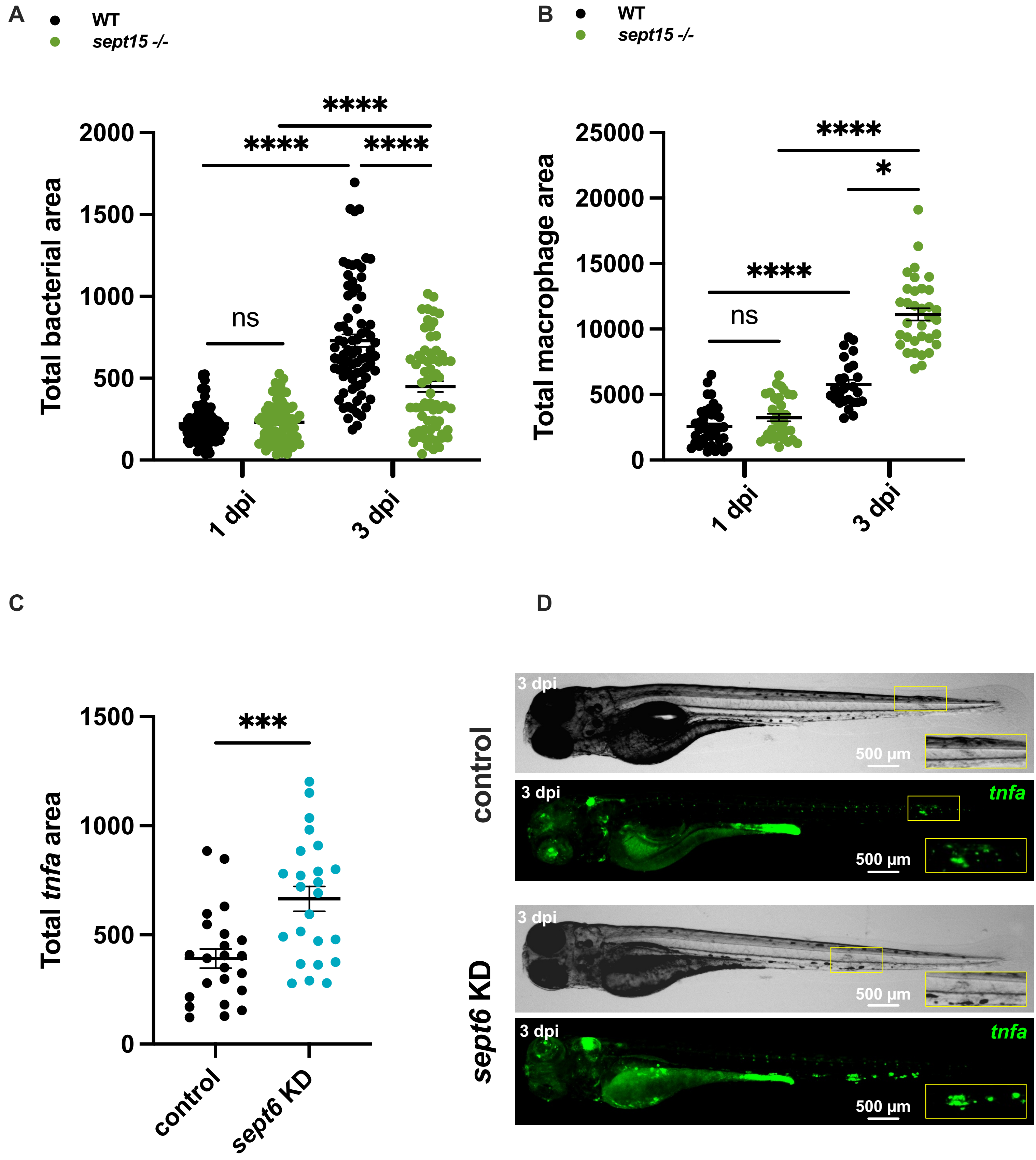

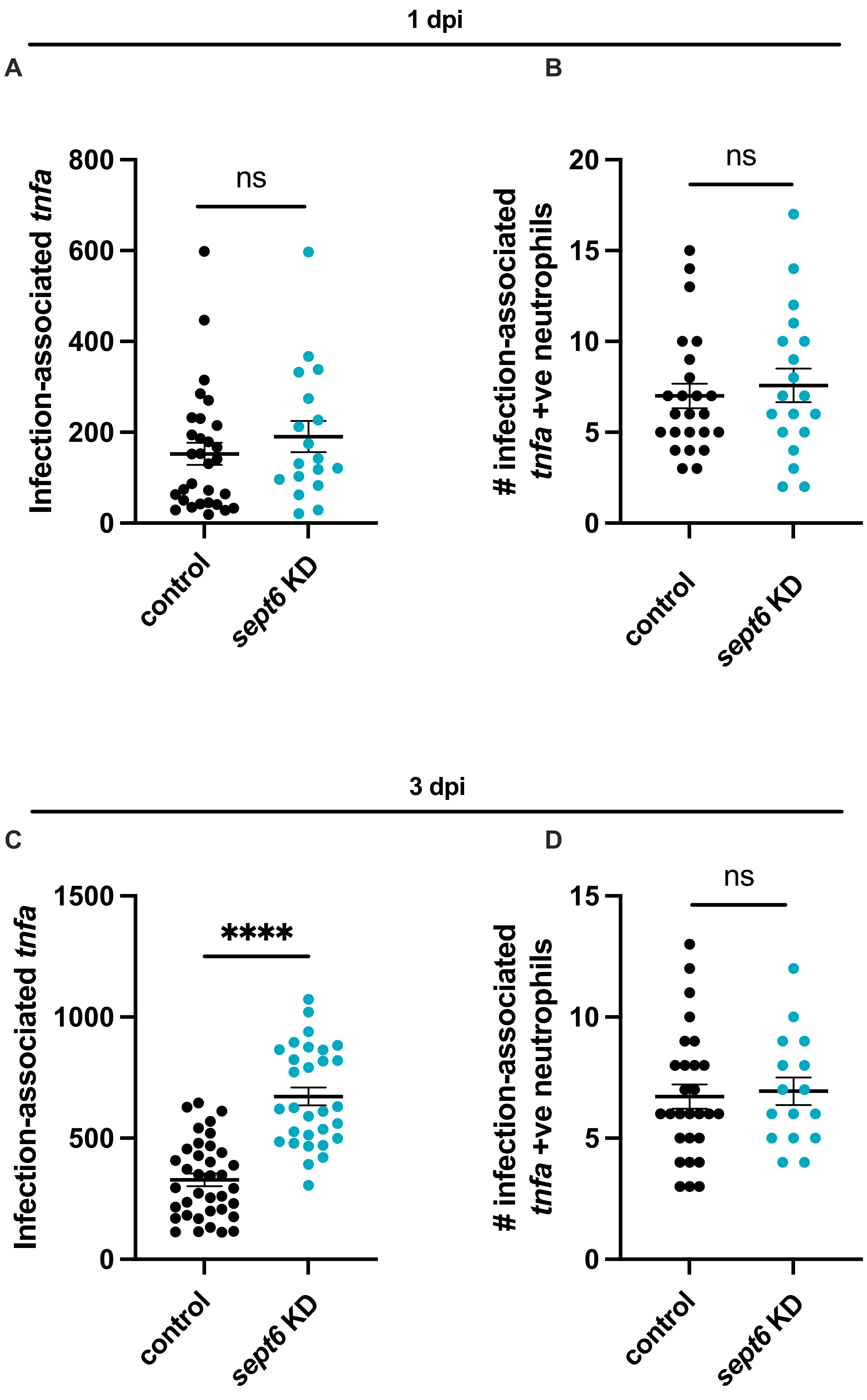

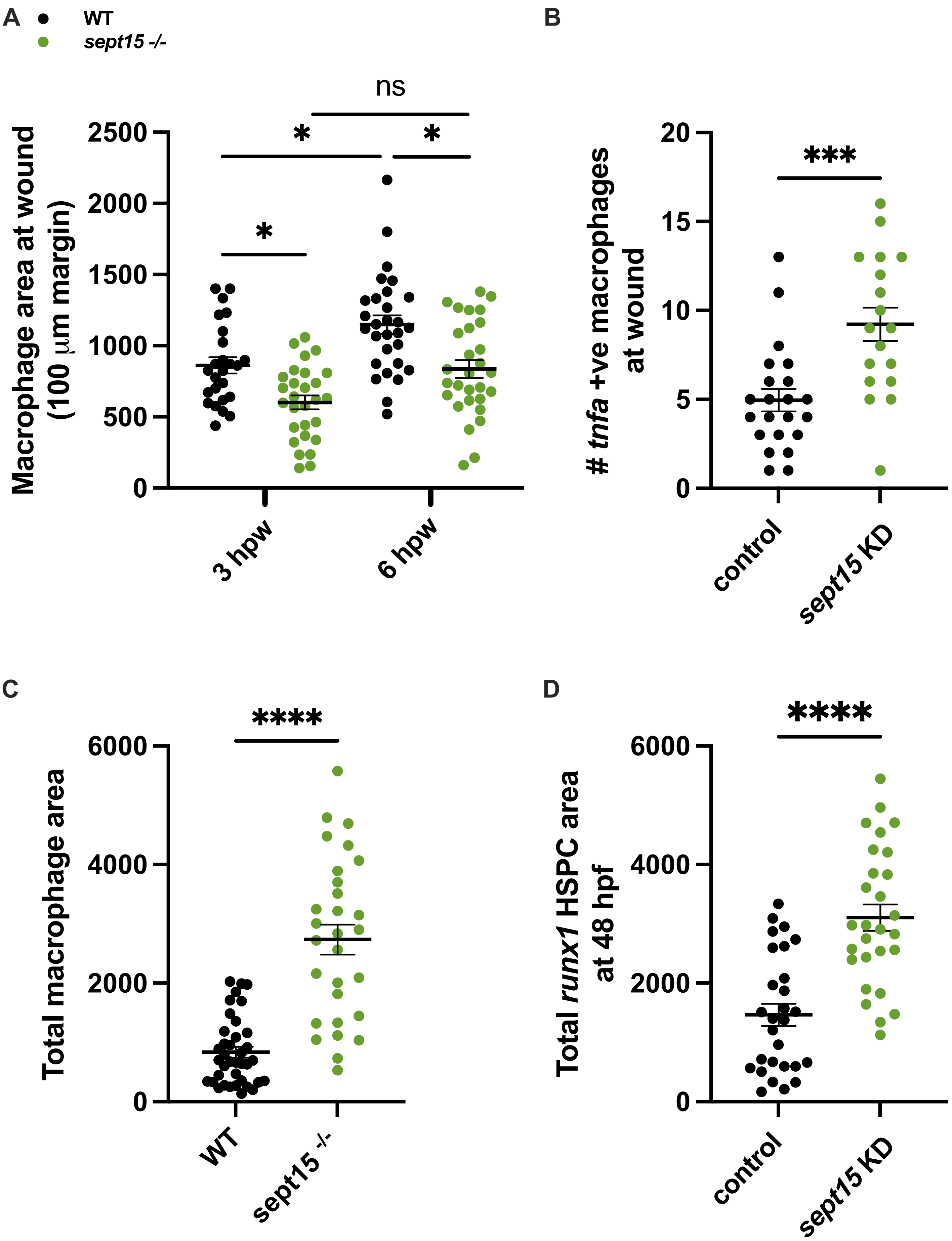

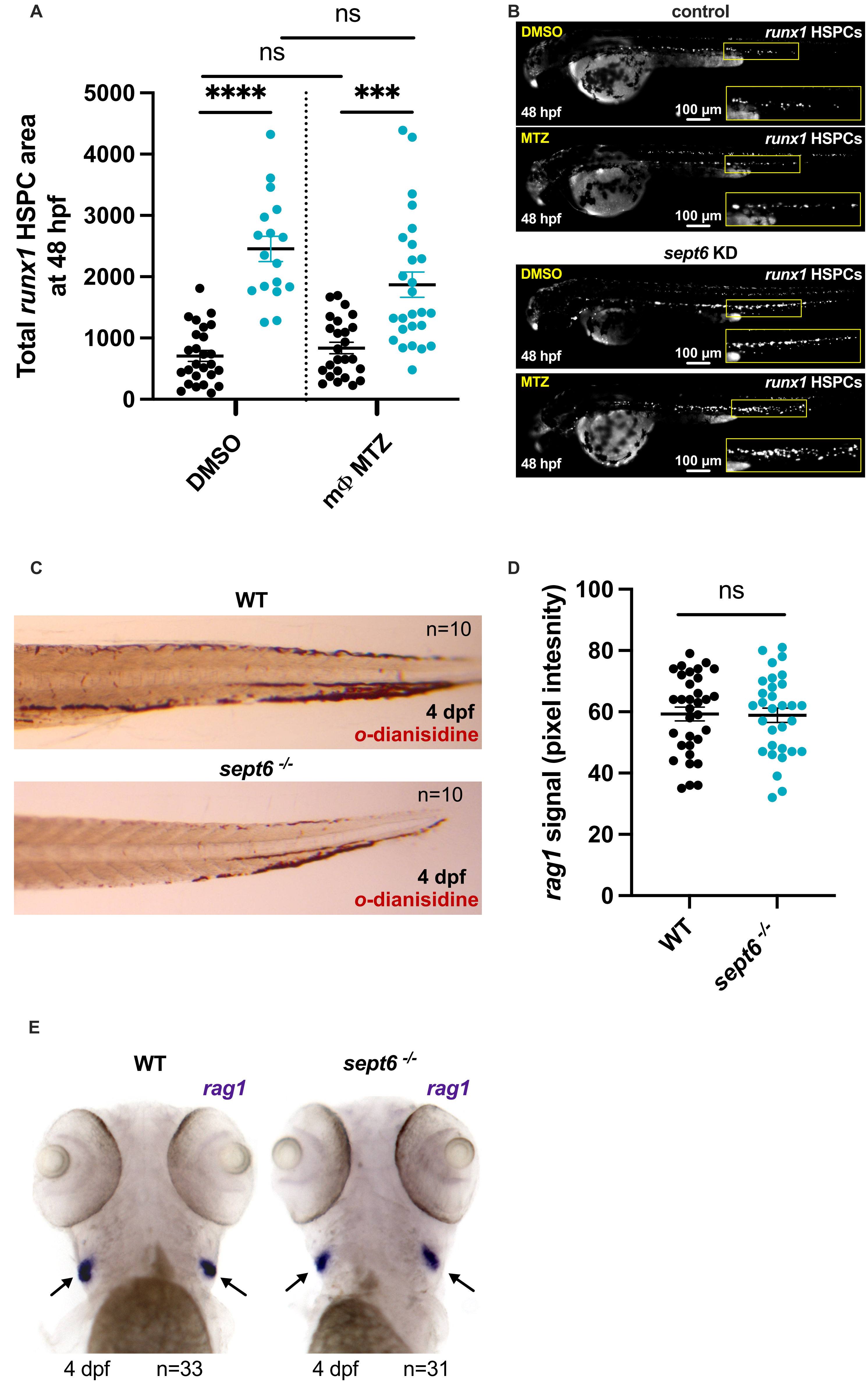
